# Hepatic Macrophage Migration Inhibitory Factor Promotes Pancreatic Cancer Liver Metastasis in NAFLD

**DOI:** 10.1101/2024.06.02.595997

**Authors:** Qian Yu, Hui Song, Liang Zhu, Xiao-ya Shi, Hai-zhen Wang, Ying-luo Wang, Rui-ning Gong, Jiu-fa Cui, Xiao-nan Yang, Ji-gang Wang, Yu Liang, Ying Chen, Xiao-wu Dong, Guo-tao Lu, Chang Li, Huan Zhang, Yan-tao Tian, Hai-tao Hu, Xin-xin Shao, Ya-bin Hu, Ashok K. Saluja, Yue Li, Ming-guang Mo, He Ren

**Author notes:** **Correspondence to He Ren**, Tumor Immunology and Cytotherapy of Medical Research Center, Center for GI Cancer Diagnosis and Treatment, the Affiliated Hospital of Qingdao University, No. 16 Jiangsu Road, Qingdao 266000, China. These authors contributed equally to this study (Qian Yu, Hui Song, Liang Zhu).

## Abstract

How pathological livers shape tumors, thereby driving pancreatic ductal adenocarcinoma (PDAC) metastasis to the liver, is poorly understood. In the present study, we focus on examining key molecules implicated in this process and assessing their translational significance. We demonstrated that patients with combined non-alcoholic fatty liver disease (NAFLD) have approximately a ninefold increased risk of developing liver metastasis compared to those without NAFLD. In mice model, NAFLD fosters an immunosuppressive microenvironment with increased tumor cell pluripotency and focal adhesion. Mechanistically, NAFLD-induced MIF mediated the progression of PDAC liver metastasis by attracting CD44 positive pancreatic cells. Hepatic MIF knockdown significantly reduced metastases burden with decreased stem-like cancer cells, tumor associated macrophages (TAMs) infiltration and focal adhesion. Targeting the MIF-CD44 axis by either a MIF tautomerase inhibitor, IPG1576, or by CD44 knockdown in tumor cells significantly attenuate liver metastasis of PDAC within the NAFLD context. Patients with PDAC liver metastasis and NAFLD had elevated hepatic MIF expression and increased number of stem-cell like cancer cells. Collectively, our study highlights a pivotal role for MIF-CD44 axis in cancer stemness and offer novel avenues for tailoring therapeutic strategies to individual patients with NAFLD as an underlying condition.

## Introduction

Pancreatic ductal adenocarcinoma (PDAC), characterized by a high metastatic burden, is currently the third most common reason for cancer deaths and projected to become the second leading cause within the next ten years[1]. The overall 5-year survival rate for patients with metastatic PDAC remains only 3% for the past two decades[2, 3]. This is in stark contrast to the encouraging progress being achieved for patients presenting with localized PDAC[3]. The primary cause of mortality in PDAC is metastasis to the liver, lungs, and peritoneal cavity[4]. Notably, almost fifty percent of individuals with PDAC are present with metastatic disease at the time of diagnosis, with majority of metastases located in the liver (76-94%)[5–7]. In addition, only about 20% of patients are diagnosed with resectable disease, and even after potentially curative resection, two-thirds of patients ultimately succumb to a distant recurrence[8, 9]. Hence, to improve patient outcomes, innovative therapeutic techniques capable of interfering with the metastatic process are required[3].

The incidence and prevalence of non-alcoholic fatty liver disease (NAFLD) are rapidly increasing worldwide. It is worth noting approximately 50% of patients undergoing pancreatic cancer surgery experience the onset of NAFLD within 12 months following the operation, while the underlying causes remain elusive[10]. In addition, according to large prospective and retrospective human datasets, individuals with NAFLD have a higher risk of liver metastasis and recurrence following resection of liver metastasis compared to patients without NAFLD[11–13]. As a result, the rising rates of NAFLD is likely to contribute to an increase in the incidence of liver metastasis, such as colorectal cancer, which has been consistently linked to NAFLD[11, 12, 14, 15].

Regarding the association between NAFLD and pancreatic cancer, limited data are available with inconsistent results of positive[16–19] or null associations[20]. Until recently, the first nationwide cohort study demonstrated an association between NAFLD and the risk of pancreatic cancer in the general population, independent of obesity[21]. However, given the prevalence of hepatic metastasis in various cancers and the presence of steatosis, we were surprised by the paucity of information regarding the relationship between NAFLD and PDAC liver metastasis. Hence, in this study, we first used clinical cohorts to reveal the correlation between hepatic metastatic PDAC and NAFLD. Owing to the limited occurrence of surgical resections in PDAC patients with liver metastasis, which makes the acquisition of liver resection surgical specimens challenging, this study primarily employed animal models for *in vivo* simulation. Among these, the murine liver metastasis model closely aligns with clinical pathological features and stands as the most frequently utilized model for liver metastasis[22]. Moreover, the driving force and key molecular features throughout this process have yet to be discovered, since the treatment approach for liver metastases may vary between individuals with or without fatty liver. In the present study, we perform a comprehensive bioinformatics analysis to characterize the cell-cell communication changes in the metastatic liver tissues, and we screened for ligand-receptor pairs with enhanced interaction between NAFLD liver and tumors: hepatocyte macrophage migration inhibitory factor (MIF) - pancreatic cancer cell surface receptor CD44.

MIF is a pleiotropic cytokine with enzymatic, chemotactic, and hormonal properties. It is stored in many kinds of cells and released by immune cells, endothelial cells, platelets, and some parenchymal cells when triggered by inflammation and stress [23]. In recent years, based on the establishment of hepatocyte-specific *Mif* knockout mouse models, studies have shown that hepatocytes are the main source of MIF in the progression of NAFLD and alcoholic liver disease (ALD) [24, 25]. In the progression of NAFLD, MIF secreted by liver cells polarizes natural killer T cells towards the pro-inflammatory and pro-fibrotic type I phenotype [24]; in ALD, MIF is an important danger signal released by liver cells in response to ethanol-induced injury [25]. Furthermore, recent studies have emphasized the pro-tumorigenic role for MIF several cancer types [26–29]. In pancreatic cancer, exosomal MIF produced from pancreatic cancer cells has been shown to have a role in liver metastasis. This is supported by the observation that exosomes lacking MIF are unable to facilitate the development of a pre-metastasis niche [29]. More recently, a highly potent small molecule inhibitor targeting the tautomerase activity of MIF, known as IPG1576, was developed, which substantially suppressed tumor growth in an orthotopic pancreatic cancer model [30]. Given the multiple roles of MIF in both NAFLD and PDAC, we hypothesize that MIF may be a key factor mediating fatty liver metastasis and a promising target for therapeutic applications.

Hyaluronan (HA) cell-surface glycoprotein receptor CD44 is a recognized marker for identifying tumorigenic and “stem-like” pancreatic cancer cells [31]. The aggressive nature of pancreatic cancer and its propensity to metastasis may be partially ascribed to a lack of effective strategies to target pancreatic stemness [32, 33] as characterized by the expression of CD44, ALDH, CD133, et al [34, 35]. Moreover, recent studies showed CD44 increases the expression and activation of β-integrin receptors, which fosters extravasation [27]. In the present study, we unraveled cancer stemness and the adhesions of metastatic cells are acquired and strengthened in the setting of NAFLD, which could be modulated by hepatic MIF knockdown or MIF inhibition. By investigating the molecular mechanisms underlying PDAC metastatic liver tumor growth in the scenario of NAFLD, we aim to seek targeted and personalized treatment strategies within this clinical context.

## METHODS

### RESOURCE AVAILABILITY

All antibodies were listed in **Data S1.**

#### Lead contact

Further information and requests for resources and reagents should be directed to and will be fulfilled by the lead contact, He Ren.

#### Patient cohorts and clinical samples

This study mainly includes cohort from Peking Union Medical College Hospital for retrospective research data from year 2013 to 2023. A retrospective analysis was conducted on computerized medical records to extract comprehensive clinical parameters (Table S1). Contrast-enhanced computed tomography and/or magnetic resonance imaging were used to assess the number and size of metastatic sites in patients with PDAC. The research excluded cases with uncommon malignant pancreatic illnesses, including as non-ductal tumors (acinar, neuroendocrine, pancreatoblastoma, and solid pseudopapillary tumors), as well as secondary malignancies (e.g., ampullary, CBD cancers)[10]. The retrospective clinical study was conducted in accordance with the Ethical Review Approval for Clinical Research Projects by the Health Commission (QYFYEC2024-50).

In research cohort II, 6 cases of PDAC liver metastasis tissues patients from the Affiliated Hospital of Qingdao University, was used for HE, multiplexed immunofluorescent staining (mIHC) and analysis. All cases of PDAC liver metastases were obtained from the Affiliated Hospital of Qingdao University from year 2018 to 2020. The demographic and clinico-pathological features were shown in Table S2. This cohort was conducted in accordance with the ethical guidelines established by the Affiliated Hospital of Qingdao University (QYFYWZLL28018).

#### Definition of NAFLD

Patients included in this study who were diagnosed with pancreatic cancer underwent abdominal computed tomographic (CT) scanning at the time of initial diagnosis in the PU cohort. The CT diagnosis involved using unenhanced CT scanning to measure the Hounsfield unit (HU) values of the liver and spleen. The liver-to-spleen (L/S) ratio of HU <1.0 was utilized for diagnosing and assessing the severity of liver fat content[36, 37]. The study excluded individuals with liver conditions like chronic viral hepatitis B/C, autoimmune hepatitis, primary biliary cholangitis, or liver cirrhosis. Additionally, those who consumed excessive alcohol (≥30 g/day for males and ≥20 g/day for females) or had a history of hepatocellular carcinoma or other liver-related malignancies were not considered for participation.

#### Mouse models of NAFLD

Wild-type (WT) mice (C57BL/6J background) were bought from Charles River Laboratories (Beijing Vital River Laboratory Animal Technology Co., Ltd, China). Mice were housed under specific pathogen-free conditions with 12-h light–dark cycle and ad libitum access to water and food in the animal core facility of the Affiliated Hospital of Qingdao University. NAFLD was induced by feeding 8-week-old mice with three different diets, including: (1) a high-fat diet (HFD) rich in saturated fat and sucrose (TP23100, Trophic Diets, Nantong, China) for 8 weeks[38]; (2) a choline-deficient, l-amino acid-defined, high-fat diet enriched with 1% cholesterol (CDAA) (TP3622657; Trophic Diets) for up to 4 weeks[39]; or (3) a methionine- and choline-deficient diet (MCD) (TP3005; Trophic Diets) for 4 weeks[40], respectively. All animal experiments adhered to protocols approved by the Research Ethics Committee of the Affiliated Hospital of Qingdao University (AHQU-MAL20231113).

#### Murine model of hepatic metastases

A preclinical, murine pancreatic tumor model of hepatic metastases model was utilized in this study which closely resembles the clinical conditions observed in patients with liver metastases[22]. Briefly, C57BL/6 male mice were injected intrasplenically with 10^6^ KPC cells in 25 µL of DMEM under isoflurane anesthesia. Mice were sacrificed on Day 15 after the initial injection.

#### Effect of IPG1576 on murine hepatic metastases model

The compound IPG1576 (30mg/kg/d) was administered twice daily via oral gavage, ensuring a minimum interval of six hours between each dose. This regimen commenced seven days prior to the establishment of the metastasis model and continued for a duration of 14 days. An equivalent quantity of solution, 5%DMSO and 95% (20% [2-Hydroxypropyl]-β-cyclodextrin), was used as vehicle for IPG1576. The recent study has detailed the absorption, distribution, metabolism, excretion, and preliminary safety analysis of IPG1576 [30].

#### Multiplex immunohistochemistry (mIHC) staining and analysis

Multiplex IHC was performed using the Opal 7-colour fluorescent IHC Kit (PerkinElmer, Massachusetts, USA) according to the manufacturer’s instruction. In brief, 4-μm thick formalin-fixed paraffin-embedded (FFPE) sections of murine or human metastatic tissues were deparaffinized and then fixed with 10% formalin fixation prior to antigen retrieval using citrate buffer (pH6) or Tris/EDTA (pH9) under high temperature and high pressure. Corresponding primary antibodies, secondary-HRP antibodies, and Opal TSA dyes were incubated with the slides for several rounds. After all sequential staining reactions, sections were counterstained with DAPI and mounted with antifade reagent (AR1109, BOSTER, Wuhan, China). The sequential multiplexed staining protocol is shown in Data S2. All mIHC slides were scanned using a Zeiss Axio Scan Z1 brightfield/fluorescence Slide Scanner at 20X magnification. For metastatic tissues obtained from patients with PDAC, whole-slide scan images were stained and analyzed. Hepatic metastatic tumor sections from mice were stained, and 3-5 representative fields (each covering an area of 14 mm²) were selected from whole-slide scan images for further analysis using HALO software (v3.6.4134, Indica Labs, USA) using Classifier (Random forest), HighPlex FL v4.2.14 module incorporated with Nearest neighbor analysis, Proximity analysis, and Infiltration analysis.

To differentiate tumor tissue from stroma, a tissue classifier leveraging the epithelial cell marker cytokeratin 19 (CK19) was utilized. The Highplex FL analysis algorithm facilitated cell phenotyping by defining nuclear detection sensitivity and minimum intensity thresholds based on staining localization (nuclear, cytoplasmic, or membrane). Sequentially, the counts of targeted cells were normalized to the total cell counts for the tumor areas to generate the density of positive cells per mm^2^ of tissue. Cell localization was assessed by measuring infiltration distances of different cell types along the tumor border. The proximity distance, representing the average number of TAMs within a 30-μm radius from the nuclear center of CD44^+^CK19^+^ tumor cells, was quantified using GraphPad Prism (v10).

#### Cell culture

Murine cell lines Kpc1199 was gifting from Professor Jing Xue (Shanghai Cancer Institute, Ren Ji Hospital, School of Medicine, Shanghai Jiao Tong University) and cultured under the recommended conditions following granted protocols, including the respective medium supplemented with 10% fetal bovine serum (FBS) and 1% antibiotics, and kept in 37 °C humidified incubators with 5% CO2. AML12 cell was supplied by the Cell Bank of Type Culture Collection of Chinese Academy of Sciences (Shanghai, China) and cultured in DMEM supplemented with 10% FBS, 1% antibiotics, 1% insulin–transferrin–selenium (Thermo Scientific), and 40ng/mL dexamethasone (Millipore, MA, USA) at 37 °C with 5% CO2 using a cell incubator.

#### Production of AAV particles

For hepatic *Mif* knockdown, three unique shRNA constructs targeting *Mif* (sh*Mif* #1, #2 and #3) and a nontargeting scrambled control (sh*Ctrl)* cloned into pLKO.1-EGFP-puro vector were used in this study. Knockdown of *Mif* was confirmed by transfection in AML12 cell followed by Western blotting. Construct #2 was then subcloned into AAV vector (AAV-U6-MCS-CMV-ZsGreen) for subsequent production of AAV particles. The AAV_sh*Ctrl* virus was used as a negative control. For hepatic *Mif* overexpression, we used the sequence of mouse *Mif* coding DNA sequence (CDS) from the National Center for Biotechnology Information (NCBI) (GenBank: NM_178398.4). CDS of *Mif* was then subcloned into the AAV vector (AAV-CMV-MCS-EF1-ZsGreen). All plasmids were confirmed by restriction enzymes and sequencing.

For the viral packaging procedure, the triple plasmid adeno-associated virus system—which consists of the pAdDeltaF6 plasmid, AAV serotype 8, and plasmid containing target gene vectors—was employed. Using HighFectin transfection reagent, the three plasmids were co-transfected with 293T cells following high purity endotoxin-free extraction of each plasmid vector. The precipitates of the cells were obtained 72 hours following transfection. Viral titre, mycoplasma, and sterility tests were conducted as part of the quality control process for viruses. The packaging, purification, and titration of the reassembled viral vector were carried out by Shanghai Integrated Biotech Solutions (Shanghai, China).

#### In vivo AAV vector treatment

AAV particles were purified, tittered, and injected through mouse with a titre of 10^11^ four weeks prior to the establishment of hepatic metastasis model.

#### SiRNA-mediated transient knockdown

The transient knockdown of *Cd44* was performed using *Cd44* siRNA (si*Cd44*) (GenePharma, Shanghai, China), which is a pool of three target-specific 19–25 nt siRNAs. KPC cells were plated in a 6-well plate at a concentration of 6×10^5^/well in a volume of [2ml culture medium. On the following day, replace the original medium with 1 ml fresh medium, and then transfected with *Cd44* siRNA or non-targeting control siRNA (si*Ctrl*) at a concentration of 150 pmol/well using GP-transfect-Mate reagent (GenePharma, Shanghai, China). The medium was changed 24 h after transfection. After 48 h of infection, observe the expression of GFP with a fluorescence microscope, and lysates were collected for western blot verification.

#### Transmigration of KPC cells toward MIF gradient

To determine the optimal concentration of MIF for migration assays, KPC cells were seeded in Transwell (8 μm pores; BIOFIL, Guangzhou, China) inserts at a density of 5 × 10^4^ cells/well in the upper chamber. The lower chamber was treated with varying concentrations of MIF (0, 50, 100 and 200 ng/mL) along with 50 μg/mL (SDF-1) serving as a positive control. All cells were harvested after 36 h, resuspended in 100 μL 1% FBS medium, and seeded in the upper Transwell inserts migration chamber, with 600 μL 1% FBS in the lower chamber. Next, we evaluate the influence of MIF on the migratory behavior of KPC cells after transfection with si*Cd44* and si*Ctrl*. Briefly, following transfection with various siRNA formulations, cells were harvested and suspended in 1% FBS medium. Subsequently, 100 μL of cells (5 × 10^4^ cells) with different treatments were added to the upper chamber. The lower chamber was filled with 1% FBS medium containing various concentrations of MIF (0 and 200 ng/mL). After incubation for 36 hours at 37°C in a 5% CO2 atmosphere, KPC cells in the upper chamber were eliminated using cotton swabs. The invaded cells crossing the membrane into lower chamber were stained with 0.1% crystal violet at 37 °C for 40min. Three randomly selected areas at a magnification of 100 × of each sample using software, and the number of migratory cells was counted.

#### Mouse liver metastases organoid culture

Mouse liver metastasis tissues were removed from the preservation solution (Kingculture™, KCX-1) and rinsed with DPBS (Thermofisher, 17105041) for three times. Tissues were cut into small pieces as much as possible with blood vessels, fat, necrosis being removed, and transferred to a clean centrifuge tube supplemented with Digestion Solution (Kingculture™, KCX) and placed in a water bath for 30 min at 37°C at 140 rpm. After digestion, the tissues were rinsed with washing solution (Kingculture™, KCQ-1) for three times. During this period, tissue fragments were blown up and down vigorously 10-20 times until no obvious cell masses appeared. The supernatant was then transferred to a new tube and centrifuged for 5 minutes at 1000 rpm. The solution was subsequently filtered through a 100 µm cell mesh sieve and centrifuged at 1000 rpm for 5 min. 4×10^5^ KPC cells and Matrigel (Corning, 356231) were mixed in a 200:1 ratio and inoculated into 6-well plates in 2 mL pancreatic cancer organoid medium (Kingculture^TM^ Organoid Growth Medium, KCW-M-8) per well. The medium was changed every 3 days.

After the organoids were constructed, they were broken down thoroughly and filtered through a 40μm filter to ensure that all cells are single cells. The diameter of the organoids should not be too large (preferably <400μm) to avoid long-term digestion, damage to cell activity, and subsequent cell sphere formation experiments.

#### Sphere Formation Assay

Liver metastasis organoids were seeded (3000 cells / well) in 96 well low-attachment culture plates in 75 uL of DEME medium (Gibco). After cells were seeded into the plates, place the cells in a 6-hour incubator for starvation treatment to consume the cell factors related to the organoid culture medium that may remain in cells. After 6 hours, add 2× pancreatic cancer tumor sphere culture medium (Kingculture™, KCW-M-13-100) in a 1:1 ratio for cultivation and observation. If cell clusters are found to be excessively aggregated, gently shake the well plate or use a wide-bore pipette tip to gently blow several times to help disperse the cells. Spheres were re-seeded up to three times every 7 days to understand the self-renewal potential, and the number of spheres (> 100 mm) for each well was evaluated after 7 and 14 days of cultivation. Each sample was seeded in 10 wells and images were taken daily for consecutive 14 days.

#### Protein extraction and immunoblotting analysis

Mice liver metastatic issue samples were harvested and homogenized in a lysis buffer (P0013, Beyotime) using a Tissue Homogenizer Low-temperature (-40[, KZ-III-FP, Servicebio). Protein lysis was separated using sodium dodecyl sulfate-polyacrylamide gel electrophoresis, transferred into a nitrocellulose membrane (Immobilon-P, Millipore), and then blocked with 5% BSA/TBST at room temperature for 1 h. The membranes were subjected to overnight incubation at 4 °C with primary antibodies. Following the washing step with TBST, the membranes were blotted using horseradish peroxidase-conjugated antibodies specific to either rabbit IgG or mouse IgG for a duration of 1 hour at room temperature. The signal was detected by the Pro e-BLOT Touch Imager using chemiluminescent substrate (WBKLS0500; Merck Millipore, Boston), and the band intensity was measured using ImageJ software (National Institutes of Health).

#### Unique molecular identifiers (UID) RNA sequencing

Total RNA was extracted from metastatic liver tumor tissues using TRIzol in accordance with the manufacturer’s instructions. The quality of the RNA was assessed by analyzing the A260/A280 ratio using the NanodropTM OneC spectrophotometer (Thermo Fisher Scientific Inc). The integrity of the RNA was verified using 1.5% agarose gel electrophoresis, followed by the quantification of qualified RNA by Qubit3.0 with QubitTM RNA Broad Range Assay kit (Life Technologies, Q10210). A total of 2 micrograms of RNA were used for building a stranded RNA sequencing library using the KC-DigitalTM Stranded mRNA Library Prep Kit for Illumina® (DR08502, Wuhan Seqhealth Co., Ltd. China)

The kit minimizes duplication bias in PCR and sequencing procedures by using a unique molecular identifier (UMI) consisting of 8 randomly generated bases to identify the pre-amplified cDNA molecules. The library products with base pair sizes ranging from 200 to 500 were subjected to enrichment, quantification, and sequencing using the DNBSEQ-T7 sequencer (MGI Tech Co., Ltd. China) with the PE150 model.

#### Histology

The tissues were fixed in a solution of 10% formalin buffered with PBS (AB1019, AOBO, China), embedded in paraffin, and cut into 4-5 µm slides. Briefly, the slides were deparaffinized with xylene and dehydrated in gradient ethanol, and the high-pressure antigen retrieval procedure was performed using the suitable antigen retrieval buffer as specified by the antibody manufacturer’s instructions. The sections were treated with a 10% goat serum blocking solution (ZLI-9021; Beijing, China) for 30 minutes at 37[, followed by incubation with corresponding primary antibodies overnight at 4[, and an anti-rabbit/mouse secondary antibody (PV-9000; Beijing, China) 1 hour at room temperature. DAB (ZLI-9018; Beijing, China) staining was performed, followed by counterstaining with hematoxylin (G1140; Solarbio, Beijing, China). Aqueous mounting material was used to attach the slides after a distilled water wash. Images were captured on a microscope (Leica) and analyzed by ImageJ software. Expression levels were scored based on staining intensity (ImageJ). To quantify the metastatic burden, H&E slides were scanned to calculate the metastatic burden with the ZYFViewer software. The area occupied by metastatic foci was then divided by the total surface area.

### QUANTIFICATION AND STATISTICAL ANALYSIS

#### Statistical analyses

Statistical analyses were conducted using GraphPad Prism software v10. The sample size is indicated in the figure legends, and details of statistical tests are provided in the respective figure legend where applicable. Appropriate tests, such as unpaired parametric Student’s t-test, ANOVA analysis, or unpaired non-parametric Mann-Whitney U test, were chosen based on the normality of data. Overall survival was estimated using the Kaplan-Meier method, and differences were assessed using the log-rank test. For infiltration and proximity analyses, two-way ANOVA followed by Šídák’s multiple comparison test was performed to identify differences between two groups at different distances. A *P*-value less than 0.05 was considered statistically significant for all studies. \**P*<0.05, ** *P*<0.01, *** *P*<0.001, and **** *P*<0.0001; “n.s.” indicates not significant.

#### Bulk RNA sequencing analysis

The sequencing data first underwent filtration using Trimmomatic (version 0.36), wherein low-quality reads were eliminated and runs containing adaptor sequences were trimmed. The Clean Reads underwent further processing using in-house scripts to remove any duplication bias that may have been introduced during library preparation and sequencing. Concisely, the clean reads were first classified into clusters based on their unique molecular identifier (UMI) sequences. The readings within the same cluster were subjected to pairwise alignment, and then, reads with a sequence similarity over 95% were isolated into a distinct sub-cluster. Once all sub-clusters were created, a multiple sequence alignment was conducted to get a single consensus sequence for each sub-cluster. Following these procedures, all mistakes and biases produced during PCR amplification or sequencing were eradicated. The UID RNA-seq experiment and high-throughput sequencing, as well as data processing and analysis, were performed by Seqhealth Technology Co., LTD (Wuhan, China). GSEA was performed using GSEA software (Broad Institute).

#### Single-cell RNA sequencing analysis

The processed data contains a large digital expression matrix with cell barcodes as rows and gene identities as columns. The processed matrix was loaded into the R with the Seurat package v4.0.5 in accordance with the introductory vignettes workflow by Seurat using their default parameters. Outlier cells were filtered using library size, number of expressed genes, and mitochondrial proportion (nFeature_RNA >200, nFeature_RNA <7500, and percent.mt<10). Public data (GSE157600) was collected from NCBI Gene Expression Omnibus to obtain the liver samples with or without tumor.53 In the outlier filtering steps, lenient criterion on mitochondrial proportion (percent.mt < 25) was applied to preserve the sample size. The counts from Cell Ranger were normalized and log2 transformed using the NormalizeData function of the Seurat package. Samples were integrated, and principal component analysis was performed by RunPCA dimensionality reduction, followed by RunUMAP. The first 20 principal components were used for clustering, with 1.0 set as the resolution parameter. In both public and our own data, some clusters were combined by gene expression similarity. The protein tyrosine phosphatase receptor type C (Ptprc, CD45) >1 parameter was used to filter our targeted myeloid cells, and macrophage UMAP was created by cell-identification criteria. Differential gene expression was assessed using the FindMarkers of the Seurat default function.

#### Data and code availability

This paper does not report original code.

Any additional information required to reanalyze the data reported in this paper is available from the lead contact upon request.

## Result

### Demographic and clinico-pathological features of PDAC patients with and without NAFLD

Between the years 2013 to 2023, a total of 1127 patients were diagnosed with PDAC at Peking Union Medical College Hospital, and 661 patients were enrolled with quantifiable liver and spleen density with their baseline characteristics are detailed in **Table 1**. Based on the results of the ratio of the liver and spleen density calculated from noncontract CT, approximately 25.72% of patients had fatty liver in all PDAC patients, which is almost the same equal to the general disease rate of NAFLD globally. However, in patients with hepatic metastasis, the rate has tremendously risen to 55.49% **(Table 1).** Furthermore, the proportion of patients with liver metastasis in those with combined NAFLD is 55.49%, whereas the proportion of patients without NAFLD who develop liver metastasis at the time of diagnosis is only 14.41% **(Fig. 1A-B**, **Table 1)**. Multivariate analyses of the prognostic factors for hepatic metastases are presented in **Table 1**. These analyses demonstrated that NAFLD was a significant factor for liver-specific progression (OR 9.12, 95% CI 5.91–14.08, *P*<.0001). Taken together, these findings suggest a close correlation between NAFLD and PDAC liver metastasis, indicating that patients with NAFLD might be more vulnerable to developing PDAC liver metastasis.

**Figure 1.**
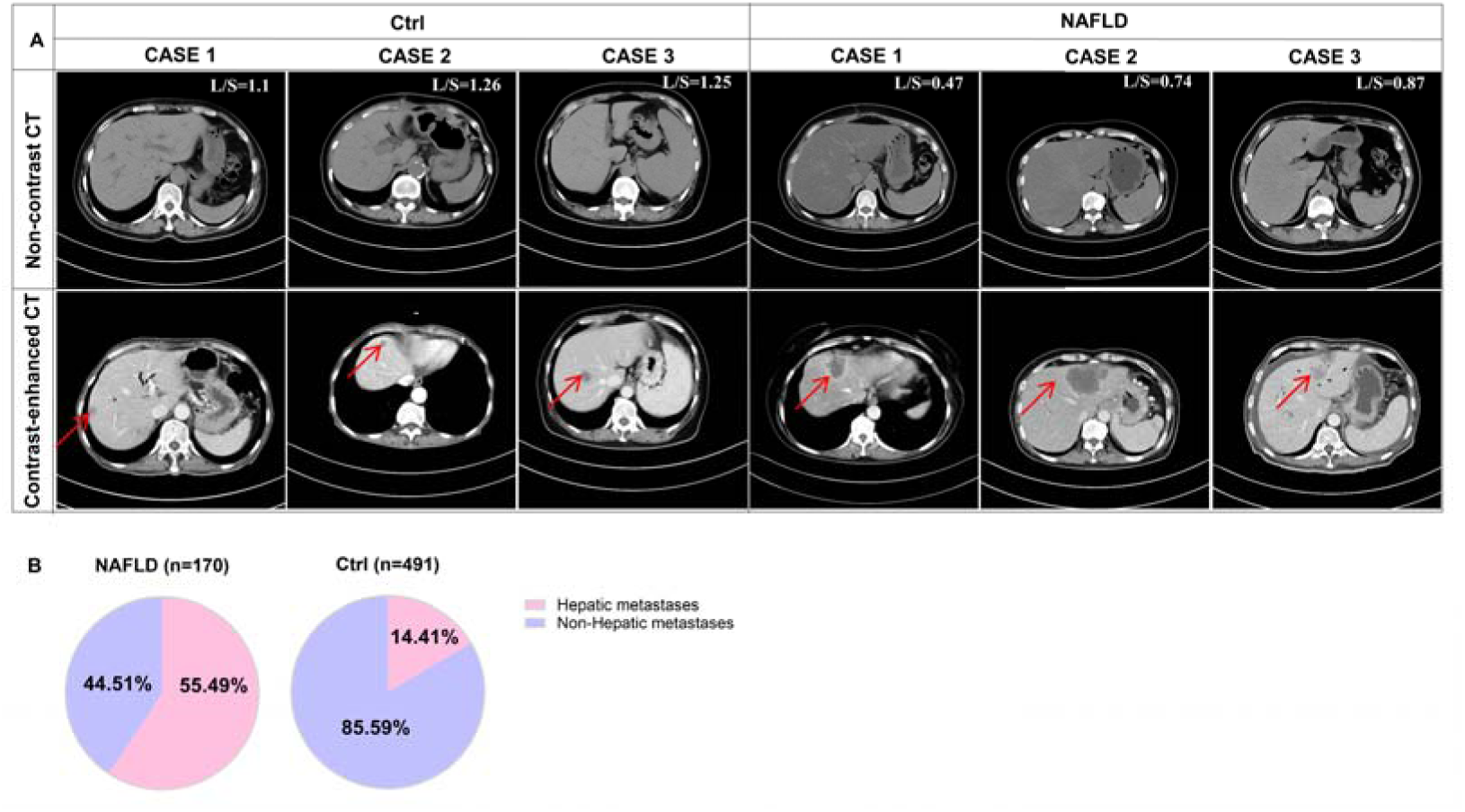
Demographic and clinico-pathological features of PDAC patients with and without NAFLD. (A) Representative non-contract and contract-enhanced CT from metastatic PDAC patients with or without NAFLD. (B) The proportion of liver metastasis in patients with pancreatic cancer diagnosed with or without concurrent NAFLD.

**Table 1.** Demographic and clinical characteristics and multivariate logistic regression analysis of PDAC liver metastases.

| Variables | Total | Control | Liver metastases | OR | P |
| --- | --- | --- | --- | --- | --- |

|  | (n = 661) | (n = 479) | (n = 182) | (CI 95%) |  |
| --- | --- | --- | --- | --- | --- |
| **Gender, n (%)** |  |  |  |  |  |
| Female | 349 (52.80) | 264 (55.11) | 85 (46.70) | 1.00 |  |
| Male | 312 (47.20) | 215 (44.89) | 97 (53.30) | 0.86<br>(0.55 ~ 1.36) | 0.5232 |
| **Age Group, year, n (%)** |  |  |  |  |  |
| < 60 | 264 (39.94) | 189 (39.46) | 75 (41.21) | 1.00 |  |
| 60~70 | 279 (42.21) | 205 (42.80) | 74 (40.66) | 1.13<br>(0.72 ~ 1.76) | 0.6019 |
| > 70 | 118 (17.85) | 85 (17.75) | 33 (18.13) | 1.51<br>(0.86 ~ 2.64) | 0.1501 |
| **Smoke, n (%)** |  |  |  |  |  |
| 0 | 509 (77.00) | 382 (79.75) | 127 (69.78) | 1.00 |  |
| 1 | 152 (23.00) | 97 (20.25) | 55 (30.22) | 1.77<br>(1.07 ~ 2.93) | 0.0258 |
| **Radical operation, n (%)** |  |  |  |  |  |
| 0 | 244 (36.91) | 138 (28.81) | 106 (58.24) | 1.00 |  |
| 1 | 417 (63.09) | 341 (71.19) | 76 (41.76) | 0.23<br>(0.15 ~ 0.35) | <.0001 |
| **NAFLD, n (%)** |  |  |  |  |  |
| 0 | 491 (74.28) | 410 (85.59) | 81 (44.51) | 1.00 |  |
| 1 | 170 (25.72) | 69 (14.41) | 101 (55.49) | 9.12<br>(5.91 ~ 14.08) | <.0001 |
| SD: standard deviation, OR: Odds Ratio, CI: Confidence Interval |  |  |  |  |  |

#### NAFLD fosters a supportive microenvironment for the metastasis of PDAC cells

We next aimed to explore the causal relationship between NAFLD and hepatic metastases in a mouse model of PDAC liver metastasis. There are several diet-induced models that are currently being used to study NAFLD, each with its own unique challenges and drawbacks[41]. The high-caloric western diet, while keeping human characteristics of metabolic syndrome, fails to mimic the complete pathology of steatohepatitis and fibrosis. On the other hand, the MCD model closely resembles human NASH histology but fails to exhibit most metabolic abnormalities such as the weight gain or glucose tolerance relevant to human NASH pathophysiology[42]. Similarly, CDAA is another commonly used model which closely replicates the pathological characteristics of clinical NAFLD [43]. Herein, we first evaluated the metastatic burden in the above mentioned NAFLD models and obtained consistent results. Regardless of the diet used, all models showed an increase in the progression of hepatic metastases **(Fig. 2A and Fig. S1A-B).** Therefore, we chose the CDAA model for the subsequent experiments. Consistent with findings from previous studies in other cancer types, such as breast and colon cancer [14, 44], significantly shortened survival was observed in CDAHFD-fed mice compared with ND-fed mice with more metastatic lesions, larger hepatic metastasis foci coverage and increased weight of livers **(Fig. 2A-C)**.

**Figure 2.**
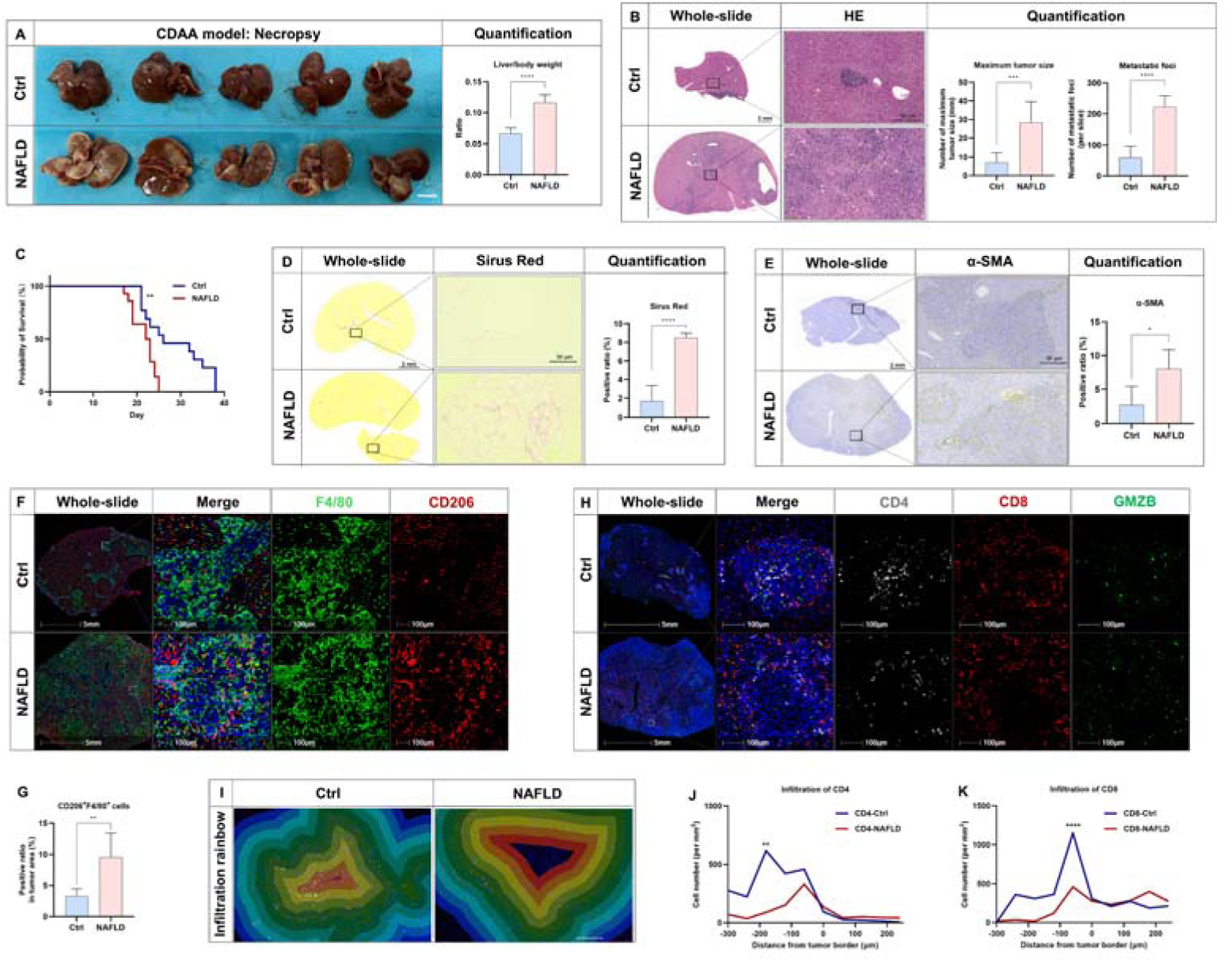
NAFLD enhances metastatic tumor growth and fosters a conducive for metastasis in the liver. After 4 weeks of NAFLD induction, mice were injected with KPC cells in the spleen and sacrificed on the 15^th^ day (n=7). (A) Representative macroscopic appearance of the liver metastases and the liver to body weight ratio in the Ctrl and NAFLD groups. (B) HE staining of liver metastases in the Ctrl and NAFLD groups with quantification of maximum tumor size and the number of metastatic foci (n=7). (C) Survival cumulative survival curves in PDAC liver metastasis mouse model stratified in the Ctrl and NAFLD groups (n=9 in Ctrl groups and n=23 in NAFLD group). (D-E) Fibrosis was assessed by Sirius Red staining and IHC staining of α-SMA using ImageJ (n=4-5). (F-G) The expression of CD206 and F4/80 were examined mIHC, and the ratio of CD206^+^F4/80^+^ cells in the tumor area was quantified by Halo software using HighPlex FL v4.2.14 module (n=4-5). (H) Immuno-suppressive microenvironment of liver metastasis was examined by mIHC staining of CD4, CD8 and GMZB. (I-K) Infiltration rainbow was shown with a band of 60μm per area and the infiltrated cell number of CD4 and CD8 per mm^2^ tissue across the tumor border was quantified (n=3). Scale bar was indicated in the individual figures. Representative pictures are shown. Data are shown as mean ± SD per group. Unpaired parametric Student’s t-test was performed to identify differences between two groups. For proximity analyses, two-way ANOVA followed by Šídák’s multiple comparison test was performed to identify differences between two groups at different distances. A *P*-value less than 0.05 was considered statistically significant. \**P*<0.05, ** *P*<0.01, *** *P*<0.001, and **** *P*<0.0001; “n.s.” indicates not significant.

Furthermore, NAFLD promoted fibrotic stroma consisting of activated cancer-associated fibroblasts (CAFs) as assessed by α-SMA, Sirius Red staining and increased angiogenesis as evident by increased CD31 expression **(Fig. 2D-E and Fig. S1C)**. Since the dynamic interactions between CAFs and immune cells orchestrates processes such as metastasis and contributes to immunosuppression in PDAC, we next evaluated the extent of immune cell infiltration and localization by mIHC. In liver metastasis tissue, antibody recognizing CD206 was used to identify the anti-inflammatory ‘M2-like’ macrophage subsets, which constituted approximately 10% of total TAM in NAFLD group versus around 5% in Ctrl group **(Fig. 2F-G)**. This also suggests that M1-like macrophages comprised most of the TAM population in the metastatic foci. In terms of tumor infiltrated leukocytes, while the proportion of CD4 memory-activated T cells did not significantly differ between the two metastatic foci groups, their spatial localization exhibited distinct patterns **(Fig. 2H-J and Fig. S1D)**. In the control group, CD4^+^ T cells were concentrated within the tumor center, whereas in the fatty liver group, they were marginalized toward the tumor edge **(Fig. 2I-J and Fig. S1D)**. Similarly, CD8^+^ T cells were even slightly more abundant in the fatty liver group, yet their primary distribution remained at the tumor edge, and their cytotoxic function was unaltered **(Fig. 2H-I and K, Fig. S1E-F)**. The exclusion of CD4 and CD8 T cells at the tumor periphery might suggest reduced immune surveillance and anti-tumor responses, and increased immunosuppression in the context of fatty liver. Taken together, fatty liver promotes PDAC liver metastasis progression in vivo and established a metastatic microenvironment characterized by increased fibrosis, enhanced angiogenesis, and immunosuppression.

#### Liver metastasis progression involves the MIF-CD44 axis triggered by NAFLD

To elucidate the underlying mechanisms through which fatty liver attract PDAC tumor cells *in vivo*, we first integrated two large open single-cell RNA-sequencing (scRNA-Seq) datasets GSE166504 which analyzes changes in individual liver cells across different stages of fatty liver, and GSE125588 to identify ductal cells isolated from PDAC tissue of 40-day-old *Kras^LSL−G12D/+^Ink4a^fl/fl^Ptf1a^Cre/+^* (*KIC*) mice. We then conducted an interactive analysis and exploration of cell-cell communication enabled by CellChat focusing on the hepatocytes and Kupffer cells which are the main functional cells and constitute approximately 75% of the total cells in the liver. We identified five targetable ligand combinations which specifically possess a kinase domain, an extracellular domain, or being secreted in the extracellular compartment **(Fig. 3A-B)**. Next, we aim to identify expressions that gradually increase with the duration of fatty liver feeding, specifically those that correlate positively with the severity of fatty liver. We found that only the expression of macrophage migration inhibitory factor (MIF) meets this screening criterion, followed by IHC validation in metastatic liver tissues in mice **(Fig. 3C-D, Fig. S2A-C)**. These observations led us to hypothesize that the upregulation of MIF in fatty liver could potentially serve as a crucial mediator in the development of NAFLD-induced liver metastasis. Subsequently, we aim to identify the specific receptors expressed on PDAC tumor cells that play a pivotal role in mediating the metastatic effects induced by MIF derived from fatty liver. Through comprehensive CellChat analysis, we have identified potential candidate receptor, CD44, known for its interaction with MIF and expression on mice KPC cell **(Fig. 3B)**. This finding was further validated through immunofluorescent staining **(Fig. 3E)**.

**Figure 3.**
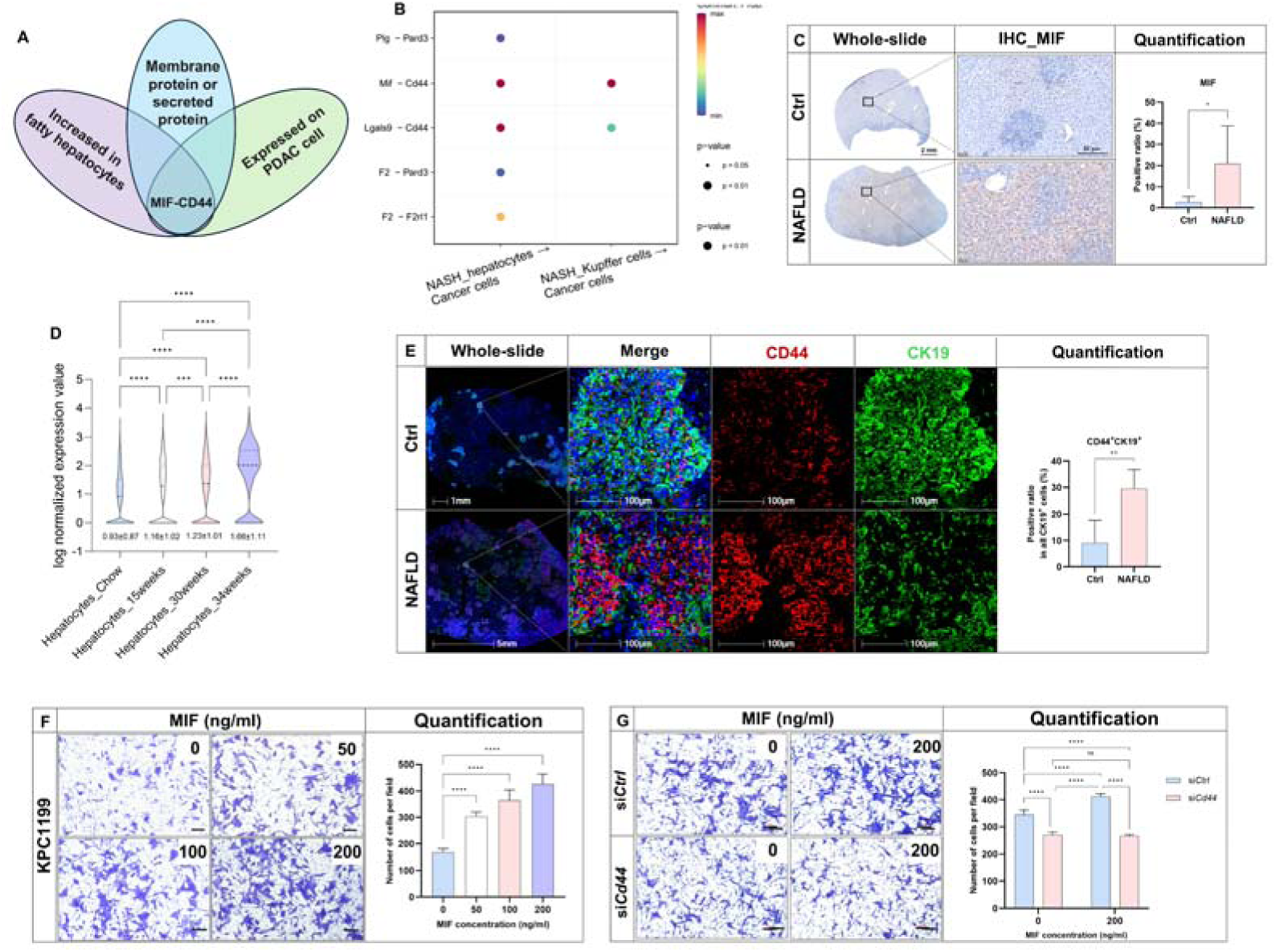
Liver metastasis progression involves the MIF-CD44 axis triggered by NAFLD. (A) Venn diagram showing screening of key molecules that mediate NAFLD-induced metastasis using single cell RNA-seq database GSE166504 and GSE125588, with the criteria of membrane protein or secreted protein. (B) A total of five ligand-receptor combinations were within the screening criteria. The ligands were produced from hepatocytes or Kupffer cells during NASH and the receptors were from pancreatic cancer cells of KIC mice. (C) Representative figures and quantification of IHC of MIF expression in the liver metastasis tissues with or without NAFLD (n=4-5/group). (D) Quantification of Mif RNA expression in the hepatocytes with different duration of high fat feeding based on GSE166504. (E) The expression of CD44 and CK19 were examined by mIHC, and the ratio of CD44^+^ cancer cells was quantified by Halo software using HighPlex FL v4.2.14 module (n=4/group). (F) MIF protein with different concentrations (0, 50, 100, 200ng/ml) was added in the lower chambers of 24-well transwell plates and 5 × 10^4^ KPC cells were seeded in the upper chambers, along with 50 μg/mL (SDF-1) serving as a positive control. (G) KPC cells were transfected with si*Ctrl* or si*Cd44* for 24 hours. MIF at a concentration of 200ng/ml was added in the lower chambers of 24-well transwell plates with 5 × 10^4^ KPC cells in the upper chambers. After incubation for 36 hours, the invaded cells crossing the membrane into lower chamber were stained with 0.1% crystal violet, and the number of migratory cells was counted. Scale bar was indicated in the individual figures. Representative pictures are shown. Data are shown as mean ± SD per group. Unpaired parametric Student’s t-test or one-way ANOVA was performed to identify differences between two groups or four groups, respectively. A *P*-value less than 0.05 was considered statistically significant. \**P*<0.05, ** *P*<0.01, *** *P*<0.001, and **** *P*<0.0001; “n.s.” indicates not significant.

CD44, a polymorphic glycoprotein, is recognized for its involvement in mediating tumor pluripotency, cell-cell adhesion and cell-matrix interactions, contributing to various cellular processes such as cellular homing, tumor invasiveness, metastasis, and angiogenesis [45]. However, the potential of hepatic MIF to function as a chemoattractant, specifically in attracting CD44^+^ tumor cells during NAFLD-induced liver metastasis, remains unknown. To address this, we first assessed the potential of mouse recombinant MIF (rMIF) at varying concentrations to stimulate the migration of KPC cells *in vitro*. The results illustrated a progressive enhancement of migration in KPC cells following treatment with increasing concentrations of rMIF **(Fig. 3F).** However, the migratory response was considerably diminished when *Cd44* was silenced **(Fig. 3G).** These data demonstrate that the chemotactic stimulation of KPC cells by MIF predominantly operates via the CD44 pathway.

To delve into the biological characteristics of metastatic pancreatic cancer within the context of fatty liver, we conducted transcriptome profiling of metastatic lesions and performed Gene Set Enrichment Analysis (GSEA) and analyzed Kyoto Encyclopedia of Genes and Genomes (KEGG) pathways based on the sequencing results. Notably, metastatic PDAC tissues associated with NAFLD exhibited increased expression of focal adhesion and proteoglycans in cancer with CD44 as a key player which significantly involved in these processes **(Data S3 and Fig. S2D-E)**. GSEA results further emphasized the involvement of CD44 in tumor pluripotency and focal adhesion pathways, supporting the aggressive nature of cancer in the presence of fatty liver **(Data S4-5 and Fig. S2F-G)**. Therefore, based on murine models, we proposed that liver metastatic foci in mice with combined fatty liver exhibit enhanced MIF-CD44 expression, which is accompanied by enhanced tumor stemness and adhesion and capabilities.

#### Hepatic MIF knockdown decreases the stemness and adhesion features of metastatic PDAC cells in the presence of NAFLD partially dependent on CD44

To investigate whether fatty liver-derived MIF promote liver metastasis in vivo, we generated liver-specific knockdown and overexpression of *Mif* using the AAV8 system, which enables precise manipulation of gene expression primarily in hepatocytes within the liver [46]. To avoid the effect of MIF on the development of NAFLD, we injected AAV8_sh*Mif* at the time of fatty liver establishment (4 weeks after CDAA feeding). Four weeks after *Mif* silencing, *Mif* mRNA was decreased; its low production level was maintained until the end of the experiment **(Fig. S3A-B, Fig. 4E)**. Metastatic liver tumor growth was significantly augmented in mice with NAFLD but was noticeably inhibited upon *Mif* silencing **(Fig. 4A-B).** A similar inhibitory effect was observed in the control group, although to a lesser extent **(Fig. 4A)**. Notably, hepatic MIF overexpression was found to promote liver metastasis only in mice fed a normal diet whereas in NAFLD mice, whereas no significant pro-metastatic effect was observed, possibly due to the high baseline MIF expression in the latter **(Fig. S4C-D).**

**Figure 4.**
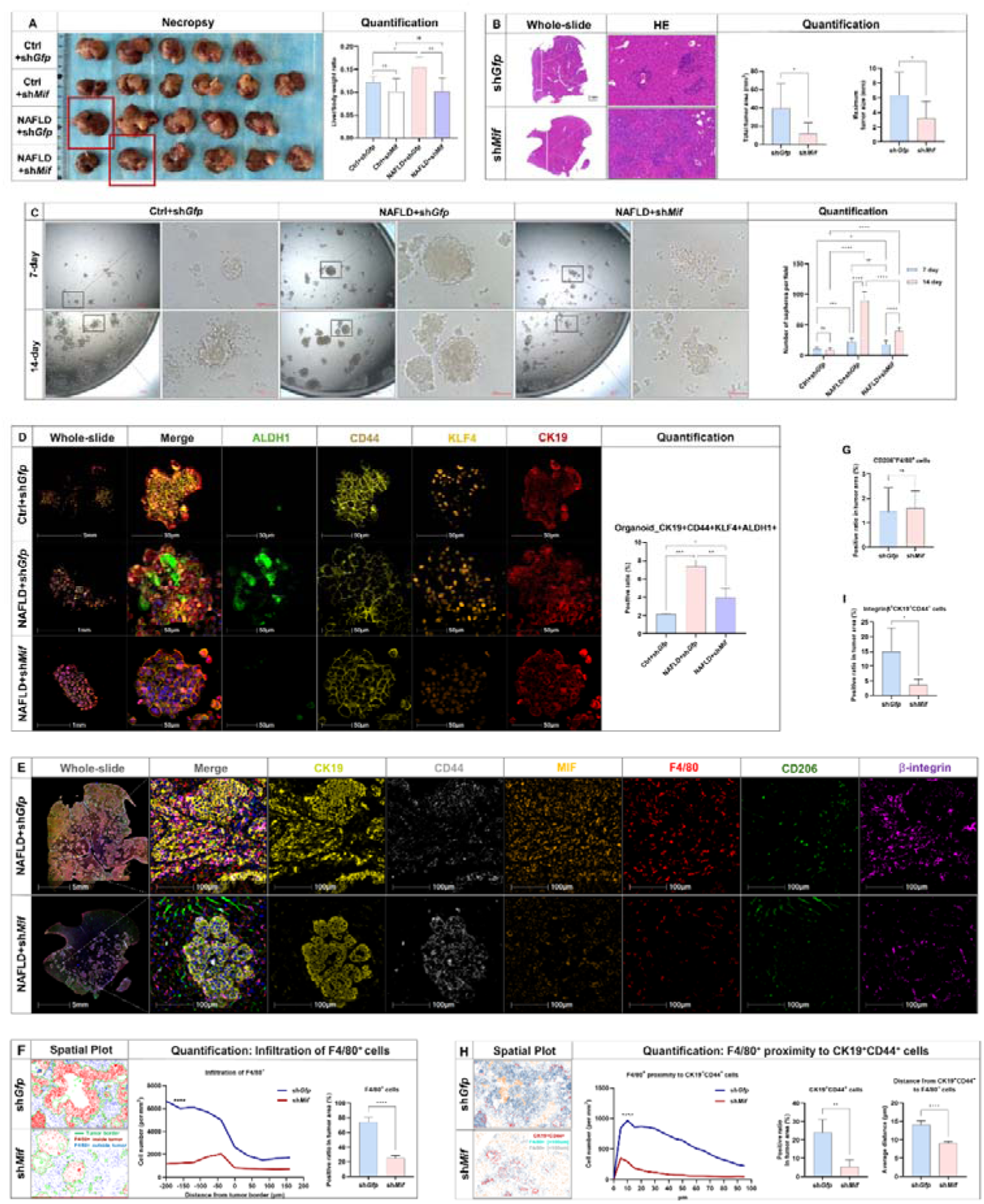
Hepatic MIF knockdown decreases the stemness and adhesion features of metastatic PDAC cells in the presence of NAFLD partially dependent on CD44. AAV_sh*Ctrl* or AAV_sh*Mif* virus were injected through the tail vein of the mice for 4 weeks to achieve hepatic MIF knockdown, followed by an ND or CDAA diet for another 4 weeks. Four groups of mice were included: (1) Ctrl+sh*Gfp*; (2) NAFLD+sh*Gfp*; (3) Ctrl+sh*Mif*; (4) NAFLD+sh*Mif*. Mice were injected with KPC cells in the spleen and sacrificed on the 15^th^ day (n=5-6). (A) Representative macroscopic appearance of the liver metastases and the liver to body weight ratio in four groups. (B) HE staining of liver metastases in the NAFLD groups with quantification of maximum tumor size and the number of metastatic foci. (C) Sphere formation assay was performed on metastatic tissues from Ctrl+sh*Gfp*, NAFLD+sh*Gfp*, and NAFLD+sh*Mif* group, respectively. 3000/well were seeded in 96-well ultra-low attachment plates in stem cell medium. 7 days later, the spheroids were resuspended and seeded again (n=2). (E) A 7-color mIHC staining including CK19, CD44, F4/80, MIF, CD206, integrinβ was performed on metastatic liver tissues from NAFLD+sh*Gfp* and NAFLD+sh*Mif* group, respectively. Representative images including whole slide, merged channels and separate channels were shown. (F-G) Spatial plot of F4/80^+^ cells were performed on metastatic liver tissues from NAFLD+sh*Gfp* and NAFLD+sh*Mif* group, respectively. Tumor border was represented in the green line, and the distribution of F4/80^+^ cells inside and outside tumor was shown in red and blue dots, respectively. The infiltrated cell number of F4/80^+^ cells per mm^2^ tissue across the tumor border was shown in the line chart, and the total number of F4/80^+^ cells and CD206^+^F4/80^+^ cells were quantified by Halo software using HighPlex FL v4.2.14 module in the column chart (n=3-5). (H) Spatial plot between CD44^+^CK19^+^ cells and F4/80^+^ cells were performed on metastatic liver tissues from NAFLD+sh*Gfp* and NAFLD+sh*Mif* group, respectively. CD44^+^CK19^+^ cells were shown as red dots. F4/80^+^ cells located more than 100 μm away from CD44^+^CK19^+^ cells appear in blue, while those within a 100-micrometer distance are depicted in gray. Proximity analysis and nearest neighbor analysis was performed on CD44^+^CK19^+^ cells and F4/80^+^ cells using Halo software (n=3-5). (I) The total number of integrinβ^+^CD44^+^ cancer cells were quantified by Halo software using HighPlex FL v4.2.14 module (n=3). Data are shown as mean ± SD per group. Unpaired parametric Student’s t-test or one-way ANOVA was performed to identify differences between two groups or among different groups, respectively. For proximity analyses, two-way ANOVA followed by Šídák’s multiple comparison test was performed to identify differences between two groups at different distances. A *P*-value less than 0.05 was considered statistically significant. \**P*<0.05, ** *P*<0.01, *** *P*<0.001, and **** *P*<0.0001; “n.s.” indicates not significant.

To ascertain the pivotal role of the CD44 pathway in mediating MIF-driven malignant traits, we conducted intrasplenic injections involving both control and *Cd44*-silenced KPC cells in CDAA-fed mice that were pre-treated with either AAV8_*Mif* or AAV8_*Gfp*. Notably, although hepatic overexpression of MIF exerted a significant enhancement on PDAC malignant characteristics, this effect was notably dampened in KPC cells where *Cd44* was silenced **(Fig. S4E-F).** Taken together, these results emphasize the central role of CD44 in orchestrating MIF-induced liver metastasis within the fatty liver microenvironment.

To establish conclusive evidence regarding the role of MIF in promoting cancer stem-like properties, we conducted experiments using tumor tissues obtained from three distinct groups: the control group, the NAFLD group, and the NAFLD group with hepatic MIF knockdown. We cultured these tissues into organoids and assessed their self-renewal abilities through spheroid formation experiments and serial replating assays. Our findings revealed significant increases in both spheroid size and number within the NAFLD group **(Fig. 4C).** Conversely, when hepatic MIF was knocked down, the typical spheroids exhibited fragmentation, and ghost-like cells were observed **(Fig. 4C)**.

Since CD44 has been identified as a reliable marker for enriching cancer stem cells (CSCs), either independently or in combination with other markers, and its multifaceted roles encompass the regulation of stemness and metastasis in various malignancies, including pancreatic cancer [47, 48], we next collected the spheroids (after second plating) and stained with stemness markers (including CD44, ALDH1, and KLF4). The results showed that the expressions of stemness markers consistently mirrored spheroid formation capabilities in above three groups **(Fig. 4D)**. These results suggest that MIF knockdown diminished stemness features of pancreatic cells in the context of fatty liver conditions. Knocking down hepatic MIF significantly reduced the overall infiltration of TAMs whilst it did not affect the proportion of M2-type TAMs **(Fig. 4E-G)**. Further analysis of the spatial distribution of TAMs within the tumor revealed a significant decrease in infiltration specifically at the central regions of the tumor **(Fig. 4F)**. On the basis of the relevant differences in the TAM and cancer cell stemness, we then conducted a spatial analysis to examine the distances of TAM in relation to CD44^+^ cancer cells. We found that hepatic MIF knockdown not only reduced the overall infiltration of TAMs and the number of CD44^+^CK19^+^ cancer cells, but TAMS existed in a more distant proximity to CD44^+^ cancer cells as compared with NAFLD group **(Fig. 4H)**. In addition to stemness signature, CD44-initiated adhesion orchestrates the expression and activation of β1-integrin receptors, promoting cancer cell attachment to endothelial cells and the extracellular matrix (ECM), thereby facilitating extravasation within the metastatic microenvironment [49]. We also found that a subset of integrinβ^+^CD44^+^ cancer cells were significantly reduced in hepatic MIF knockdown group **(Fig. 4I)**. In summary, these results indicate that hepatic MIF knockdown may result in reduced cancer stemness and adhesion in NAFLD, which was partially dependent on CD44.

#### Therapeutic targeting of MIF-CD44 axis inhibits metastatic tumor growth

Subsequently, we evaluated the therapeutic effects of intervening in the MIF-CD44 axis. Notably, IPG1094 stands as the only FDA-approved small molecule inhibitor targeting MIF. Moreover, IPG1576, representing the third generation of such compounds, offers enhanced properties in comparison to IPG1094 **(Data S6)**. We thus assessed the therapeutic effects of targeting hepatic metastasis using IPG1576 in the context of NAFLD. The results showed that IPG1576 administration significantly led to reduced liver metastasis as read out by the necropsy and HE quantification **(Fig. 5A-B)**. Consistent with results of hepatic *Mif* knockdown, MIF inhibitor significantly reduced the number of TAMs in the center and the border of the tumor without changing the ratio of M1 and M2 TAMs **(Fig. 5C-G)**. In addition, MIF inhibition in NAFLD led to a reduction in integrinβ^+^CD44^+^ cancer cells **(Fig. 5C and H).** Furthermore, spatial analysis revealed there was more distant proximity between TAMs and CD44^+^ cancer cells upon MIF inhibition **(Fig. 5I-J)**. In terms of cancer cell stemness, MIF inhibition in NAFLD led to a reduction in CD44^+^ or ALDH^+^ or CD44^+^/KLF4^+^ cancer cells, followed by Western blotting analysis showing that stemness and adhesion markers were all reduced as demonstrated by CD44, OCT3/4, SOX2, Nanog, KLF4, integrin V, and integrinβ **(Fig. K-O)**. On the other hand, after knocking down CD44 expression in KPC cells, we injected them into the spleen to establish liver metastasis model. Importantly, compared to the control group, the progression of liver metastasis was significantly attenuated as we observed a reduction in tumor area and a decrease in liver weight **(Fig. P-Q)**. These findings suggest that targeting MIF-CD44 axis within the context of fatty liver background may have therapeutic effects.

**Figure 5.**
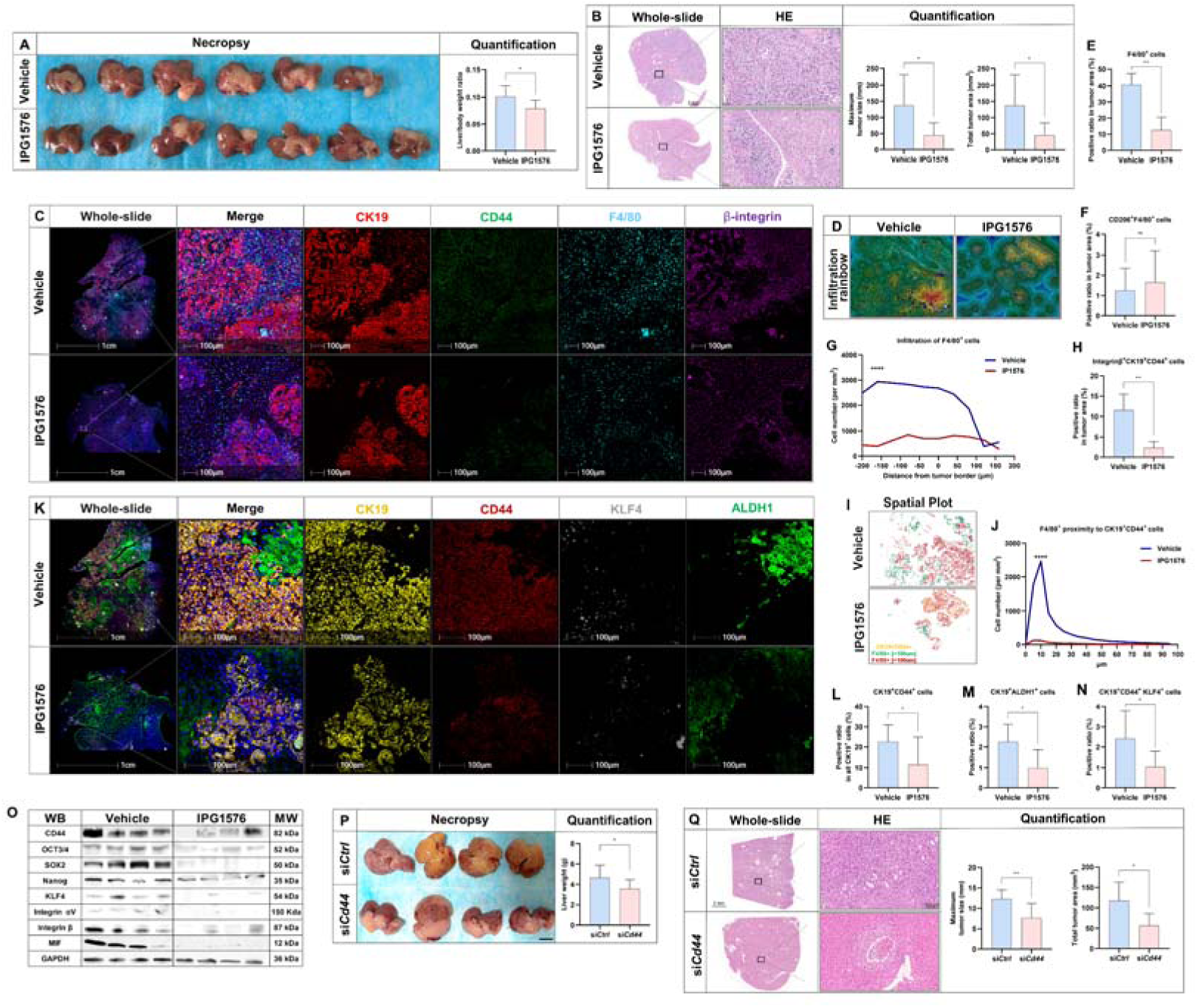
Therapeutic targeting of MIF-CD44 axis inhibits metastatic tumor growth. Mice were fed with a CDAA diet for 4 weeks before the establishment of the liver metastasis model. The compound IPG1576 (30mg/kg/d) was administered twice daily via oral gavage and commenced seven days prior to the establishment of the metastasis model and continued for a duration of 14 days. An equivalent quantity of solution, 5%DMSO and 95% (20% [2-Hydroxypropyl]-β-cyclodextrin), was used as vehicle for IPG1576 (n=6-7). (A) Representative macroscopic appearance of the liver metastases and the liver to body weight ratio in four groups. (B) HE staining of liver metastases in the NAFLD groups with quantification of maximum tumor size and the total area of metastatic foci. (C) A 4-color mIHC staining including CK19, CD44, F4/80, and integrinβ was performed on metastatic liver tissues from vehicle and IPG1576 group, respectively. Representative images including whole slide, merged channels and separate channels were shown. (D) Infiltration rainbow was shown with a band of 60μm per area. (E-F) The ratio of F4/80^+^ cells and CD206^+^F4/80^+^ cells in tumor area were quantified using Halo software. (G) The number of F4/80^+^ cells per mm^2^ tissue across the tumor border was quantified based on the infiltration rainbow (Figure 5D). (H) The total number of integrinβ^+^CD44^+^ cancer cells were quantified by Halo software using HighPlex FL v4.2.14 module. (I-J) Spatial plot between CD44^+^CK19^+^ cells and F4/80^+^ cells were performed on metastatic liver tissues from vehicle and IPG1576 group, respectively. CD44^+^CK19^+^ cells were shown as yellow dots. F4/80^+^ cells located more than 100 μm away from CD44^+^CK19^+^ cells appear in green, while those within a 100-micrometer distance are depicted in red. Proximity analysis was performed on CD44^+^CK19^+^ cells and F4/80^+^ cells using Halo software. (K) A 4-color mIHC staining including CK19, CD44, KLF4 and ALDH1 was performed on metastatic liver tissues from vehicle and IPG1576 group, respectively. Representative images including whole slide, merged channels and separate channels were shown. (L-N) The quantification of stem-like cancer cells which are positive for CD44, ALDH, or CD44/KLF4 by Halo software using HighPlex FL v4.2.14 module. (O) WB gels showing the expression of stem-like markers from metastatic liver tissues from vehicle and IPG1576 group, respectively. (P) Mice were fed with a CDAA diet for 4 weeks. KPC cells were transfected with si*Ctrl* of si*Cd44* for 24h prior to the establishment of liver metastasis model. Representative macroscopic appearance of the liver metastases and the liver to body weight ratio in two groups (n=4). (Q) HE staining of liver metastases in the si*Ctrl* and si*Cd44* groups with quantification of maximum tumor size and the total area of metastatic foci (n=4). Scale bar was indicated in the individual figures. Representative pictures are shown. Data are shown as mean ± SD per group. Unpaired parametric Student’s t-test was performed to identify differences between two groups. For proximity analyses, two-way ANOVA followed by Šídák’s multiple comparison test was performed to identify differences between two groups at different distances. A *P*-value less than 0.05 was considered statistically significant. \**P*<0.05, ** *P*<0.01, *** *P*<0.001, and **** *P*<0.0001; “n.s.” indicates not significant.

#### MIF-CD44 expression pattern in PDAC patients with NAFLD

Given the infrequent surgical resection for PDAC patients with liver metastases and that obtaining large liver tissue samples is challenging, we collected a limited number of samples from liver oligometastases in individuals with pancreatic cancer. These samples were then subjected to IHC and mIHC validation and spatial analysis. Consistently, in liver tissue from patients with liver metastases and concurrent fatty liver, we observed significantly elevated expression of MIF, particularly within the non-tumor areas, as well as CD44 positive tumor cells **(Fig. 6A-C)**. Additionally, mIHC staining revealed an increase M2-type TAMs positive for CD163 and CD68 within the tumor regions, whereas CD206 was poorly expressed, indicating that CD163 was a better marker in this scenario **(Fig. 6D-E)**. Notably, compared to the control group, the number of total TAMs were not significantly different but were found to be in closer proximity to CD44^+^ tumor cells with a stem-like phenotype **(Fig. 6F)**. These findings corroborate previous results obtained in animal models, highlighting the specific expression pattern of the MIF-CD44 axis and its associated spatial distribution in liver metastasis patients with concurrent fatty liver, making it a promising target for tailored interventions in the context of PDAC, suggesting that inhibition of MIF-CD44 axis has translational potential.

**Figure 6.**
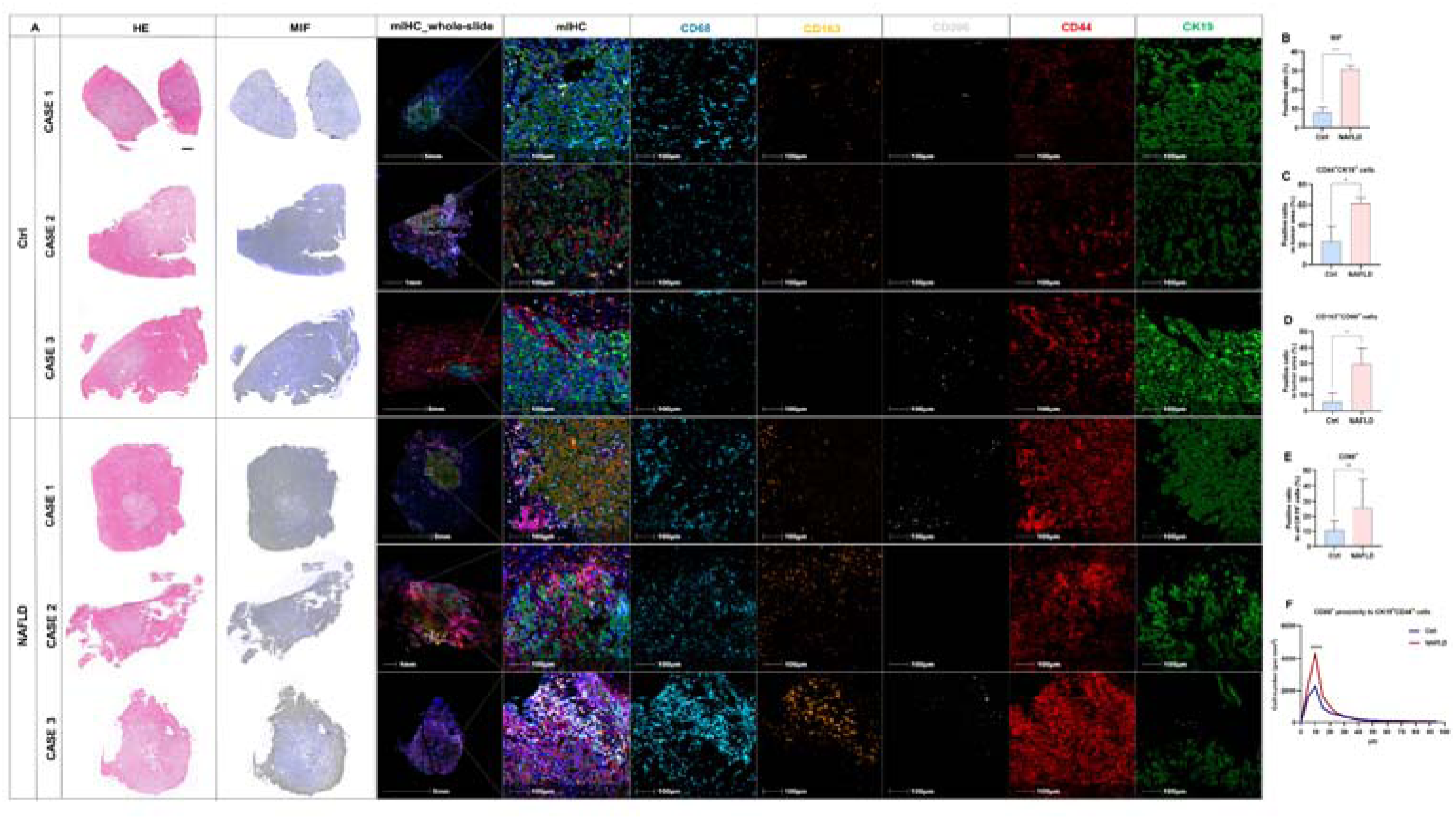
MIF-CD44 expression pattern in PDAC patients with NAFLD. (A) Metastatic PDAC tissues from patients were stained with HE, MIF by IHC, and a 5-color panel including CD68, CD163, CD206, CD44, CK19 by mIHC (n=3 for Ctrl group and NAFLD, respectively). (B) The expression of MIF was assessed by Image J based on whole-slide scanning and quantification. (C-E) The ratio of CK19^+^CD44^+^ cells, CD163^+^CD68^+^ cells and CD68^+^ cells were quantified by Halo software using HighPlex FL v4.2.14 module based on whole-slide scanning. (H) Proximity analysis between CD44^+^CK19^+^ cells and CD68^+^ cells were performed on the whole slide of metastatic liver tissues from PDAC patients using Halo software. Data are shown as mean ± SD per group. Unpaired parametric Student’s t-test was performed to identify differences between two groups. For proximity analyses, two-way ANOVA followed by Šídák’s multiple comparison test was performed to identify differences between two groups at different distances. A *P*-value less than 0.05 was considered statistically significant. \**P*<0.05, ** *P*<0.01, *** *P*<0.001, and **** *P*<0.0001; “n.s.” indicates not significant.

## Discussion

When it comes to tumor patients, the coexistence of different liver diseases does not significantly alter treatment plans or the selection of medications. Two key factors might contribute to this: Firstly, the intricate relationship between various pathological liver conditions and liver metastasis in pancreatic cancer remains poorly understood. Secondly, there has been a lack of comprehensive foundational research to identify specific targets.

Our study zeroes in on the clinical connection between NAFLD and liver metastasis in pancreatic cancer. We have established that pancreatic cancer patients with concurrent NAFLD face a higher likelihood of developing metastasis. Animal models further validate the cause-and-effect relationship between fatty liver and liver metastasis. Building upon this model, we have pinpointed enhanced biological features related to tumor cell stemness and adhesion within the context of fatty liver. Furthermore, we have discovered MIF, a critical factor secreted by fatty liver, which directs pancreatic cancer cells through the CD44 pathway. Targeting the MIF-CD44 axis effectively curbs liver metastasis progression with coexisting fatty liver in vivo. Thus, our research offers novel insights and lays the groundwork for personalized treatment strategies in pancreatic cancer patients with concurrent liver conditions.

In colorectal cancer, numerous clinical studies have consistently demonstrated that patients with fatty liver exhibit significantly higher rates of tumor incidence and recurrence compared to those without fatty liver, and multiple basic research studies have confirmed this causal relationship and delved into the underlying mechanisms [11, 14, 50, 51]. Consistent results have also been observed in primary cancers of liver, breast cancer, prostate, and melanoma [15, 16]. As previously mentioned, clinical studies yield contradictory conclusions regarding the link between fatty liver and pancreatic cancer or liver metastasis, and there is a scarcity of basic research articles on this subject, which can be attributed to several factors. Firstly, precise fatty liver diagnosis often relies on magnetic resonance imaging (MRI) or ultrasound. However, tumor patients typically undergo contrast-enhanced MRI or CT scans, while non-contrast CT scans are available in only a limited number of hospitals. Consequently, identifying an adequate number of pancreatic cancer patients with concurrent fatty liver for research purposes poses a challenge. Secondly, patients with pancreatic cancer liver metastasis rarely undergo surgical resection, which makes obtaining liver samples for clear fatty liver diagnosis challenging. In our research, we addressed this limitation by enrolling patients from three clinical centers based on data from MR, ultrasound, or non-contrast computed tomography (CT) scans. Our findings revealed that approximately 55.49% of liver metastasis patients had concurrent fatty liver, surpassing the average prevalence of 25%. Interestingly, in colorectal cancer, this ratio is approximately 50%[51], suggesting that patients with fatty liver are more susceptible to metastasis.

Recent investigations into steatotic liver inducing the tumor metastasis have yielded intriguing findings[51, 52]. Notably, the Seki team revealed that extracellular vesicles released by fatty liver hepatocytes, containing microRNAs, play a critical role in driving tumor cell migration[51]. These vesicles inhibit LATS2, leading to increased YAP transcription factor activity within tumor cells. Consequently, these tumor cells upregulate CYR61 expression, fostering an immunosuppressive microenvironment dominated by M2 macrophages and reduced CD8 T cells. This study underscores the importance of considering the specific microenvironment of metastatic sites, beyond the tumor itself, aligning with the “seed and soil” theory. Our study findings support this hypothesis and demonstrated that fatty liver-secreted factor, MIF, may drive the malignant transformation of pancreatic cancer liver metastasis. MIF has been identified as an important component of tumor-derived exosomes facilitating tumorigenesis and metastasis. Specifically, in pancreatic cancer, research led by Lyden, et al. highlighted MIF-containing exosomes released by pancreatic cancer cells play a crucial role in liver metastasis by establishing pre-metastatic niches [53]. In addition, a recent study further demonstrated the functional domains of MIF protein including tautomerase and TPOR activities are required for the inhibition of MDSC differentiation induced by pancreatic cancer–derived exosomes[30]. Accordingly, an MIF tautomerase inhibitor called IPG1576 was developed, which effectively suppressed pancreatic cancer progression by inhibiting MDSC differentiation and enhancing CD8^+^ T cell infiltration. Our study also employs this inhibitor and proposes that liver-derived MIF promotes the progression of fatty liver combined with liver metastasis. Targeting MIF effectively mitigates this progression, further expanding the indications for IPG1576. However, whether MIF is also secreted in the form of exosomes remain unclear, warranting further exploration.

Current evidence demonstrated that the pro-tumorigenic effect of MIF primarily occurs through interaction with the CD74/CD44 complex, which activates the downstream pathways, e.g. ERK1/2, Src, Akt, etc., particularly in the context of immune cell regulation [54, 55]. Our original research for the first time revealed that MIF protein can attract pancreatic cancer KPC cells through chemotaxis based on in vitro experiments. Notably, this chemotactic response is dependent on the presence of CD44 receptors expressed by KPC tumor cells. However, the exact MIF domain responsible for this activity and its potential reliance on CD74 necessitate further exploration.

In our research, we examined the spatial relationship between TAMs and CD44^+^ tumor cells exhibiting stem-like properties. Notably, downregulating hepatic MIF not only decreased the TAM population but also influenced their distribution toward CD44 positive tumor cells. The interactions between cancer stem cells (CSCs) and immune cells are most evident in the context of TAMs[56]. CSCs can recruit monocytes through chemokines like CCL2 and CSF1[57] Simultaneously, they exert a regulatory influence on TAMs, nudging them toward polarization to bolster tumor growth [58, 59]. Reciprocally, TAMs can modulate CSC self-renewal via factors such as IL-6 and TGF-beta[60, 61]. These interactions involve receptors on the surface of tumor stem cells, including PTPRZ1 and EPHA4, although other receptors remain to be elucidated[56, 62, 63]. Our study highlights that MIF knockdown or inhibition in fatty liver significantly increases the distance between TAMs and CSCs while reducing their density around CSCs. Unraveling whether the MIF-CD44 axis orchestrates cellular communication between TAMs and CSCs beckons further exploration.

In summary, our study identified the clinical relevance between NAFLD and PDAC metastasis through retrospective cohort studies across three centers. Fatty liver-induced MIF promotes an immunosuppressive tumor microenvironment with increased TAM infiltration, tumor cell stemness and adhesion in the context of fatty liver, which is partially dependent on CD44 on pancreatic tumor cells. Targeting the MIF-CD44 axis effectively ameliorates the progression of liver metastasis combined with fatty liver in vivo. Overall, our study provides theoretical support for precision medicine for pancreatic cancer patients with liver metastasis.

### Limitations of study

Our study highlights that MIF secreted by the fatty liver may enhance pancreatic cancer liver metastasis via the CD44 receptor. However, we have not specifically identified the dominant cell type within fatty liver responsible for MIF secretion. Although some studies suggest hepatocyte-rather than Kupffer cell-derived MIF play a primary role in the NALFD and ALD, our research has not specified this aspect further. Furthermore, we have not quantified or assessed the functionality of immune cell subpopulations in the liver’s immune microenvironment, which is a direction for our next steps. Lastly, since our study primarily focuses on the metastatic organ (the liver), we have not fully simulated the tumor’s transition from the arterial to venous system using in situ or KPC models. Thus, further studies are needed to overcome these limitations.

## Acknowledgement

We would like to thank the research laboratory directed by Professor Wang Yibing, Jinan, Shandong, the First Afiliated Hospital of Shandong First Medical University, Shandong Provincial Qianfoshan Hospital, for their supporting on experimental instruments. This work was partly supported by National Natural Science Foundation of China (82303933, 82125026, 82330081) and Natural Science Foundation of Shandong Province (ZR2022QH241). The authors declare no competing interests in this study.

