## Supplementary figures for "Hepatic Macrophage Migration Inhibitory Factor Promotes Pancreatic Cancer Liver Metastasis in NAFLD"

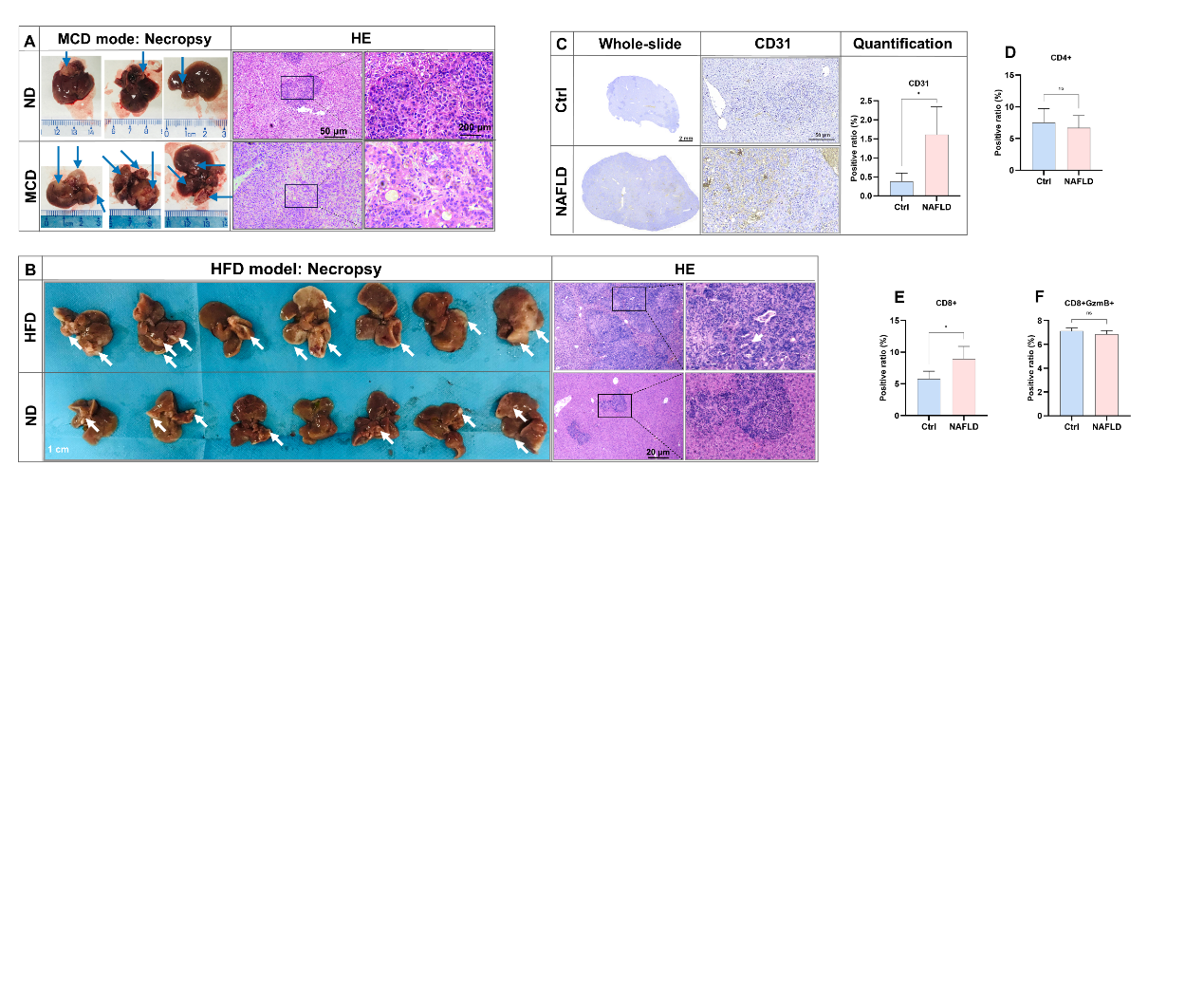
**Supplementary figures**

**Figure S1. NAFLD enhances metastatic tumor growth in the liver.** (A) After 4 weeks of MCD induction, mice were injected with KPC cells in the spleen and sacrificed on the 15th day (n=7). Representative macroscopic appearance of the liver metastases and HE staining. (B) After 8 weeks of HFD induction, mice were injected with KPC cells in the spleen and sacrificed on the 15th day (n=7). Representative macroscopic appearance of the liver metastases and HE staining. (C) Angiogenesis was assessed by IHC staining of CD31 with quantification in CDAA model (n=4-5). (D-F) The quantification of CD4, CD8 and GMZB were examined mIHC and quantified by Halo software using HighPlex FL v4.2.14 module (n=4-5).

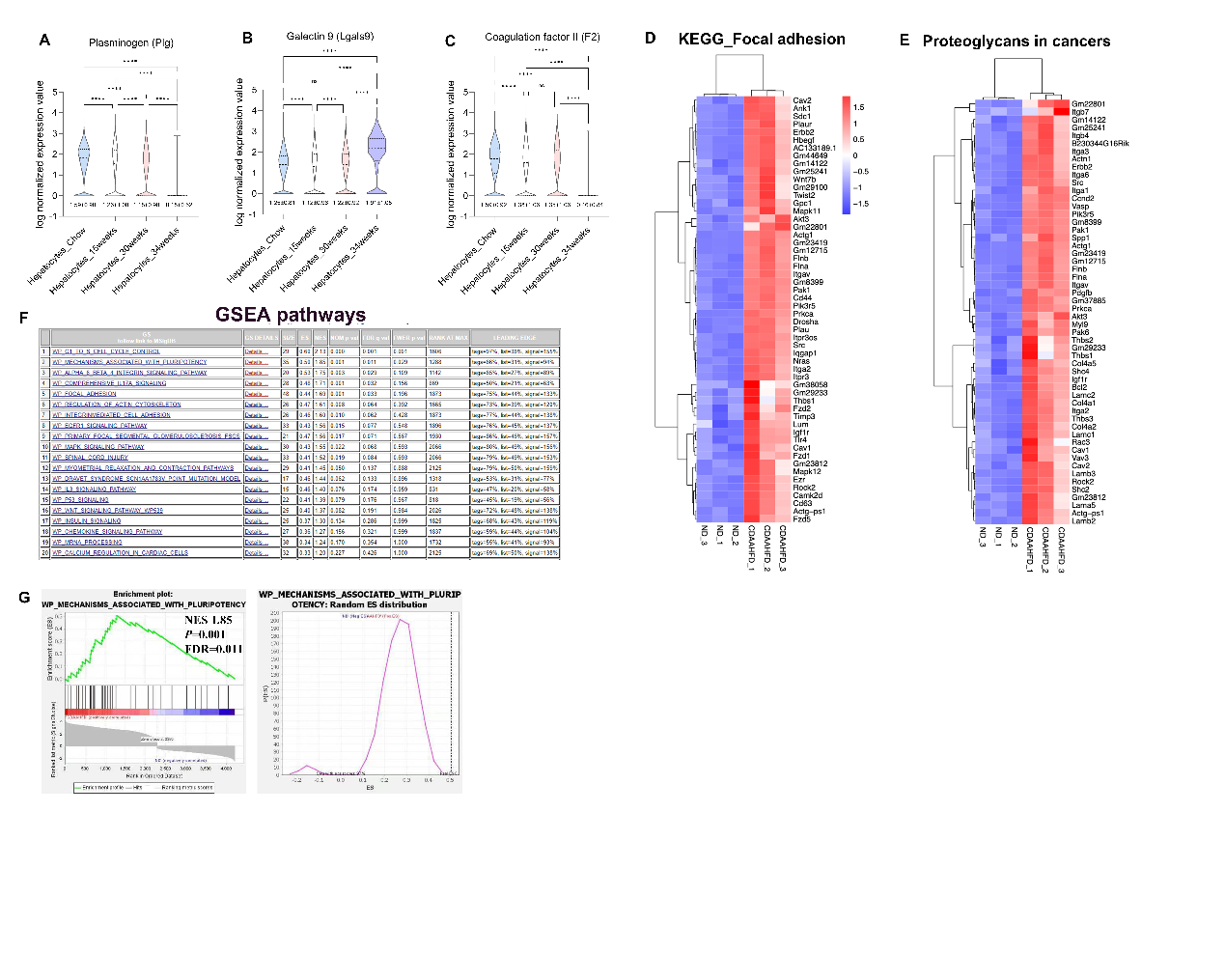

**Figure S2. Liver metastasis progression involves the MIF-CD44 axis triggered by NAFLD.** (A-C) Quantification of RNA expressions of *Plg*, *Lgals9* and *F2* in the hepatocytes with different duration of high fat feeding based on GSE166504. (D-E) Heatmap of the top three upregulated KEGG pathways (Focal adhesion and Proteoglycans) based on transcriptomic analysis of liver metastatic tissues (n=3 per group) (CDAA model). (F) Top 20 GSEA pathways involved in the NAFLD-induced metastasis based on transcriptomic analysis of liver metastatic tissues (n=3 per group) (CDAAmodel). (G) RNA-seq for tumor samples from Figure 1A (n = 3/group). GSEA for gene sets associated with tumor pluripotency gene sets in metastatic liver tissues from ND-fed and CDAA-fed mice. NES, normalized enrichment score; FDR, false discovery rate.

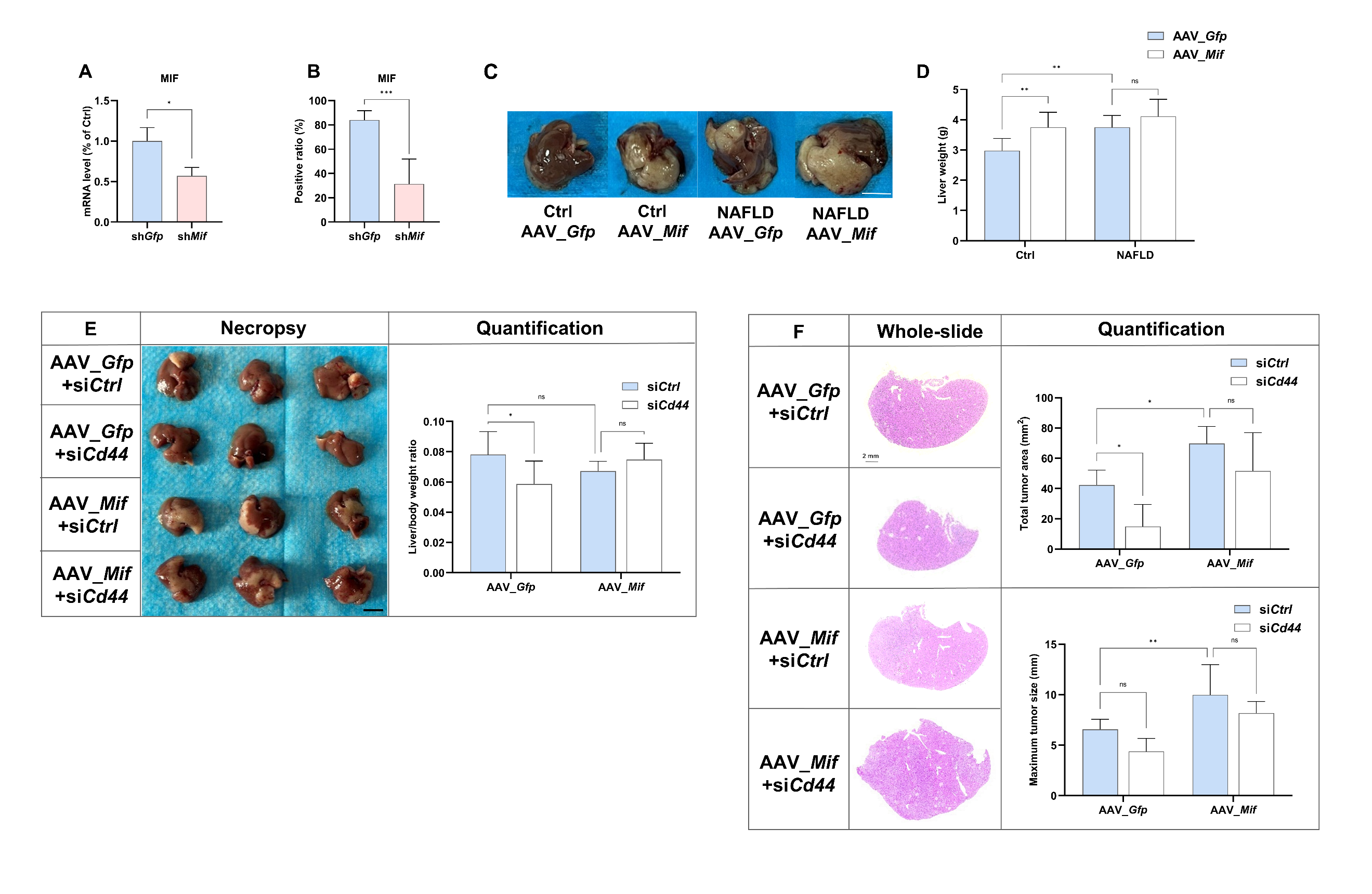

**Figure S3. CD44 is the critical factor for MIF-mediated liver metastasis in NAFLD** (A) After 4 weeks of CDAA induction, AAV-sh*Gfp* or AAV_sh*Mif* particles were injected through the tail vein to achieve hepatic MIF knockdown. Liver tissues were harvested for RNA extraction after 4 weeks of virus injection and examined. Hepatic mRNA levels of *Mif* were examined by qPCR (n=4-5/group). (B) MIF expression was examined by mIHC and quantified by Halo software corresponding to Figure 4E. (C-D) After 4 weeks of CDAA induction, AAV-*Gfp* or AAV_*Mif* particles were injected through the tail vein to achieve hepatic overexpression of MIF. Mice were injected with KPC cells in the spleen after four weeks of virus injection and sacrificed on the 15th day (n=6). Representative macroscopic appearance of the liver metastases was shown, and liver weight of each group was quantified. (E-F) 8-week mice were fed with a normal diet, and AAV-*Gfp* or AAV_*Mif* particles were injected through the tail vein to achieve hepatic overexpression of MIF. KPC cells were transfected with si*Ctrl* or si*Cd44* for 24 hours before injected in the spleen of the mice (n=6-9). Representative macroscopic appearance of the liver metastases and the liver to body weight ratio in four group (E). HE staining of liver metastases in four groups and quantification of maximum tumor size and the total area of metastatic foci (F).
